# Caspase-2 regulates S-phase cell cycle events to protect from DNA damage accumulation independent of apoptosis

**DOI:** 10.1101/2021.03.30.437768

**Authors:** Ashley Boice, Raj K Pandita, Karla Lopez, Melissa J Parsons, Chloe I Charendoff, Vijay Charaka, Alexandre F. Carisey, Tej K. Pandita, Lisa Bouchier-Hayes

## Abstract

In addition to its classical role in apoptosis, accumulating evidence suggests that caspase-2 has non-apoptotic functions, including regulation of cell division. Loss of caspase-2 is known to increase proliferation rates but how caspase-2 is regulating this process is currently unclear. We show that caspase-2 is activated in dividing cells in G1- and early S-phase. In the absence of caspase-2, cells exhibit numerous S-phase defects including delayed exit from S-phase, S-phase-associated chromosomal aberrations, and increased DNA damage following S-phase arrest. In addition, caspase-2-deficient cells have a higher frequency of stalled replication forks, decreased DNA fiber length, and impeded progression of DNA replication tracts. This indicates that caspase-2 reduces replication stress and promotes replication fork protection to maintain genomic stability. These functions are independent of the pro-apoptotic function of caspase-2 because blocking caspase-2-induced cell death had no effect on cell division or DNA damage-induced cell cycle arrest. Thus, our data supports a model where caspase-2 regulates cell cycle events to protect from the accumulation of DNA damage independently of its pro-apoptotic function.

## Introduction

Caspase-2 is a member of the caspase family of proteases that have essential roles in the initiation and execution of apoptosis.^1^ Caspase-2 has been implicated in apoptosis in response TO a variety of cell stressors including: DNA damage, heat shock, metabolic stress, and endoplasmic reticulum (ER) stress.^2^ However, the requirement for caspase-2 for cell death in each of these contexts has been subject to debate.^3^ Despite this, caspase-2 has been shown to function as a tumor suppressor in several murine models. Caspase-2-deficient mice show accelerated tumorigenesis in murine models of hematological cancers (Eμ-*Myc*-driven lymphoma^4^ and *Atm* knockout-associated lymphoma^5^), and solid tumors (*Kras*-driven lung tumors^6^ and MMTV-*c-neu*-driven mammary tumors^7^). Caspase-2-deficient tumors from these mice often showed increased features of genomic instability, including aneuploidy,^5^ and cell cycle defects, such as bizarre mitoses and an increased mitotic index.^7^ Interestingly, caspase-2-deficient tumors have shown minimal differences in apoptosis compared to wild type tumors^5, 6^ suggesting that, in addition to inducing apoptosis, caspase-2 may carry out its tumor suppressive function by regulating other cellular functions such as cell cycle.

Although it has been postulated that caspase-2 plays a role in cell cycle arrest or cell cycle checkpoint regulation,^4^ the exact phase of the cell cycle where caspase-2 primarily functions remains unclear. Caspase-2 knockout cells proliferate at higher rates.^4, 8^ In addition, caspase-2 deficiency is associated with impaired cell cycle arrest in response to DNA damage.^4^ The activation platform for caspase-2 is a large molecular weight protein complex called the PIDDosome, comprising of the proteins PIDD and RAIDD.^9^ PIDD overexpression induces growth suppression that dependent on RAIDD and partially dependent on caspase-2.^4, 10^ PIDD is a p53 target gene and PIDD can induce growth arrest in cells that are wild type for p53 but not in cells where p53 is absent or mutated.^11^ Caspase-2 induces p53-dependent cell cycle arrest in response to supernumerary centrosomes resulting in MDM2 cleavage and p53 stabilization. Because these studies place caspase-2 upstream of p53, it is possible that caspase-2 can also regulate cell cycle in a p53-independent manner. This has been noted for caspase-2 dependent apoptosis that can be either p53-dependent^12–14^ or p53-independent.^15, 16^ Caspase-2 has been shown to be activated by several different inducers of DNA damage including etoposide, cisplatin, and camptothecin.^13, 17–19^ However, apoptosis can often proceed in the absence of caspase-2 in response to these triggers and, when apoptosis is reduced, it is rarely completely blocked by the absence of caspase-2.^17, 18, 20^ In particular, while caspase-2 is efficiently activated by topoisomerase I inhibitors such as camptothecin, death in response to camptothecin is only reduced by around 50% in caspase-2-deficient cells.^17^ Of note, these drugs are also potent inducers of cell cycle arrest.^21, 22^ Inhibition of topoisomerase I triggers both cell cycle arrest and apoptosis following stalling of DNA replication forks.^23^ Fork stalling and fork collapse results in single strand DNA breaks that, in the absence of repair, are converted to double-strand breaks (DSBs), serving as a trigger for cell death or cell cycle arrest.^24^

Here, we demonstrate that in response to replication stress caspase-2 plays a key role in protecting from stalled replication forks and the subsequent DNA damage. Caspase-2 is activated during G1 and early S-phase and loss of caspase-2 is associated with several additional S-phase-related cell cycle defects including S-phase specific chromosomal aberrations and delayed exit from S-phase following arrest. In addition, we show that these defects in cell cycle regulated events are independent of caspase-2’s ability to induce apoptosis.

## Materials and Methods

### Chemicals and antibodies

The following antibodies were used: anti-Caspase-2 (clone 11B4 from Millipore); anti-phospho-ATM (Ser1981) (clone 10H11 from Invitrogen and clone 10H11.E12 from Cell Signaling Technology), anti-ATM (clone D2E2 from Cell Signaling Technology), anti-phospho-ATR (Ser428) (polyclonal from Cell Signaling Technology), anti-phospho-ATR (Thr1989) (polyclonal from Cell Signaling Technology), anti-ATR (clone E1S3S from Cell Signaling Technology), anti-phospho-Chk1 (Ser317) (clone D12H3 from Cell Signaling Technology), anti-Chk1 (clone 2G1D5 from Cell Signaling Technology), anti-phospho-Chk2 (Thr68) (clone C13C1 from Cell Signaling Technology), anti-Chk2 (polyclonal from Cell Signaling Technology), anti-caspase-3 (polyclonal from Cell Signaling Technology), anti-actin (C4 from MP Biomedicals), γH2AX (EMD Millipore). All cell culture media reagents were purchased from Thermo Fisher (Carlsbad, CA, USA). Unless otherwise indicated, all other reagents were purchased from Sigma-Aldrich (St. Louis, MO, USA).

### Cell culture and cell lines

HeLa cells were grown in Dulbecco’s Modified Essential Medium (DMEM) containing FCS (10% (v/v)), L-glutamine (2 mM), and Penicillin/ Streptomycin (50 I.U./50 μg/ml). Mouse embryonic fibroblasts (MEF) were grown in the same medium supplemented with sodium pyruvate (1 mM), 1X non-essential amino acids, and beta-mercaptoethanol (55 μM). Litter-matched *Casp2^+/+^* and *Casp2^−/−^* MEF were generated and transduced with E1A and Ras as previously described.^19^ Briefly, early passage MEF were simultaneously transduced with frozen supernatants of the retroviral expression vectors pBabePuro.H-ras (G12V) and pWZLH.E1A (provided by S.W. Lowe and G. Hannon). After 48 hours, the cells were harvested by mild trypsinization, seeded at 1 × 10^5^ cells/well in 6 well plates and cultured for 10 days in media containing 0.5 μg/ml of puromycin and 40 μg/ml of hygromycin for selection of the transduced viruses. HeLa.C2-Pro BiFC cells were generated as previously described.^17^ HeLa.C2-Pro BiFC cells expressing Am-Cyan-Geminin and *Casp2^+/+^* and *Casp2^−/−^* MEF stably expressing vector or BcLXL were generated by retroviral transduction. Gryphon-Ampho packaging cells were transiently transfected with Am-Cyan-Geminin (pRetroX-SG2M-Cyan Vector from Takara), pLZRS or pLZRS.BCLxL using Lipofectamine 2000 transfection reagent (Invitrogen, Grand Island, NY, USA) according to manufacturer’s instructions. After 48 hours, virus-containing supernatants were cleared by centrifugation and incubated with *Casp2^+/+^* and *Casp2^−/−^* MEF followed by selection in neomycin for pRetroX or zeocin for pLZRS vectors.

### CRISPR/Cas9 gene editing

*Casp2* was deleted from U2OS and HeLa cells using an adaptation of the protocol described in ref. 25.^25^ Protospacer sequences for each target gene were identified using the CRISPRscan scoring algorithm (www.crisprscan.org (Moreno-Mateos et al., 2015)). DNA templates for sgRNAs were made by PCR using pX459 plasmid containing the sgRNA scaffold sequence and using the following primers:

ΔCasp2(76) sequence: ttaatacgactcactataGGCGTGGGCAGTCTCATCTTgttttagagctagaaatagc
ΔCasp2(73) sequence: ttaatacgactcactataGGTGTGGAGGGCGCCATCTAgttttagagctagaaatagc
universal reverse primer: AGCACCGACTCGGTGCCACT.

sgRNAs were generated by *in vitro* transcription using the Hiscribe T7 high yield RNA synthesis kit (New England Biolabs). Purified sgRNA (0.5 μg) was incubated with Cas9 protein (1 μg, PNA Bio) for 10 min at room temperature. HeLa or U2OS cells were electroporated with the sgRNA/Cas9 complex using the Neon transfection system (Thermo Fisher Scientific) at 1005 V, 35 ms, and two pulses. Knockout was confirmed by PCR and western blot.

### Microscopy

Cells were imaged using one of two microscope systems. The first is a spinning disk confocal microscope (Carl Zeiss MicroImaging, Thornwood, NY, USA), equipped with a CSU-X1A 5000 spinning disk unit (Yokogawa Electric Corporation, Japan), multi laser module with wavelengths of 458 nm, 488 nm, 514 nm, and 561 nm, and an Axio Observer Z1 motorized inverted microscope equipped with a precision motorized XY stage (Zeiss). Images were acquired with a Zeiss Plan-Neofluar 40x 1.3 NA or 64x 1.4 NA objective on an Orca R2 CCD camera using Zen 2012 software (Zeiss). The second is a Leica SP8-based Gated Continuous Wave laser scanning confocal microscope (Leica Microsystems) equipped with a white-light laser, operated by Leica software. Cells were plated on dishes containing glass coverslips coated with fibronectin (Mattek Corp. Ashland, MA, USA) 24 h prior to treatment. For time-lapse experiments, media on the cells was supplemented with HEPES (20mM) and 2-mercaptoethanol (55μM). Cells were allowed to equilibrate to 37°C in 5% CO_2_ prior to focusing on the cells in an incubation chamber set at 37°C. For the Leica system, samples were sequentially excited by a resonant scanner at 12,000Hz using a white light laser (excitation wavelengths: 470nm, 514nm and 587nm) and emitted light was collected through a HC PL APO 63x/1.47 oil objective and through a pinhole of 1.5AU to be finally quantified by 3 HyD detectors with the following spectral gates: 475-509nm / 520-581nm / 593-702nm and during an acquisition window set from 0.7 to 6 ns using time gating.

### Immunofluorescence

Cells plated on dishes containing glass coverslips coated with fibronectin were washed in 3 × 2 ml PBS and fixed in 2% (w/v) paraformaldehyde in PBS pH 7.2 for 10 min. Cells were washed for 3 × 5 min in PBS followed by permeabilization in PBS, 0.15% (v/v) Triton for 10 min. Cells were blocked in FX image enhancer (Invitrogen) for 30 min and then stained with the anti-γH2AX antibody at a 1:100 dilution in PBS, 2% (w/v) BSA for 1 h. After washing in PBS, 2% (w/v) BSA, the cells were incubated with anti-mouse Alexa Fluor 555-conjugated secondary antibody (Invitrogen) at a 1:500 dilution in 2% (w/v) BSA for 45 min. Cells were washed in PBS, stained with SYTO™ 13 green fluorescent nucleic acid stain (400 nM, Thermo Fisher) in PBS and incubated at room temperature in the dark for 1 h prior to imaging

### Image analysis

At least 30 individual images were acquired for each treatment. For quantitation of nuclei, cells were stained with NucBlue™ Live ReadyProbes™ stain (Thermo Fisher) and individual nuclei were counted using the python-based CellProfiler™ software (Massachusetts Institute of Technology, Broad Institute of MIT and Harvard). Greyscale TIFF images were corrected for illumination differences using an illumination function. Images were smoothed by Gaussian method followed by separation using Otsu with an adaptive thresholding strategy. Cells were then declumped by shape and automatically counted. For quantitation of γH2AX foci, the number of foci per cell was calculated from RGB TIFF images using FoCo, a graphical user interface that uses Matlab and ImageJ.^26^ In the nuclei channel, Huang thresholding was used to separate cells. A Top-Hat transformation was used on foci images for noise reduction. Foci were segmented by using the threshold found by Otsu’s method as minimal peak height in H-maxima transform. Foci per cell were then calculated automatically.

For analysis of time-lapse imaging, cells expressing the BiFC components were identified by fluorescence of the linked mCherry protein in stable cell lines. The raw signal from mCherry was first improved using Noise2Void1 algorithm for denoising using deep learning approach. The Noise2Void 2D model was trained from scratch for 100 epochs on 148480 image patches (image dimensions: (520,520), patch size: (64,64)) with a batch size of 128 and a mse loss function, using the Noise2Void 2D ZeroCostDL4Mic2 notebook (v 1.12) on Colab cloud services (Google) and then applied to all images from the mCherry channel. Next, using a same platform (Google Colab) and a beta version of ZeroCostDL4Mic notebook2, the CellPose algorithm3 was applied to this “enhanced” mCherry channel in order to create accurate masks of each cell (Object diameter: 20um, Flow threshold 0.7 and Cell probability threshold −0.7). Raw data and CellPose masks were imported into Aivia 9.8 (Leica Microsystems) and the mean fluorescence intensity was measured in each of the Cyan and Venus channels using the CellPose-derived masks as regions of interest. After manual verification and curation of the lineage attributions, data was exported for further analysis in MATLAB (r2021a, MathWorks) where all the timelines from the different cells were aligned according to their cytokinesis time point, and were scaled by the following formula: scaled point = (Max − *x*)/MaxDifference, where Max equals the maximum value in the series, *x* equals the point of interest, and MaxDifference equals the maximum minus the minimum value in the series.

### Flow cytometry

For cell cycle analysis, cells were treated as indicated (see figure legends). Treatment media was exchanged for media containing BrdU (10 μM). After 30 min cells were collected by trypsinization, fixed, permeabilized, and stained with BrdU-FITC antibody and 7-AAD using the BD Pharmingen™ FITC BrdU flow kit according to the manufacturer’s protocol. Briefly, the cells were fixed and permeabilized with BD Cytofix/Cytoperm buffer for 15 min at room temperature, then with the secondary permeabilization buffer, BD Cytoperm Permeabilization Buffer Plus, for 10 min on ice and finally with BD Cytofix/Cytoperm buffer for 5 min at room temperature. Cells were washed in BD Perm/Wash buffer and centrifuged at 300 × g between each step. To uncover the BrdU epitope, the cells were treated with DNAse (30 μg per 10^6^ cells) for 1 h at 37°C, washed in BD Perm/Wash buffer, and centrifuged at 300 × g. The cells were then incubated in a 1:50 dilution of FITC-labeled anti-BrdU antibody in BD Perm/Wash buffer for 20 min at room temperature, washed and centrifuged at 300 × g. The cells were then resuspended in Stain Buffer (FBS) containing 10 μl of 7-aminoactinomycin D (7-AAD)/10^6^ cells. Cells were quantitated for BrdU and 7-AAD positivity by flow cytometry. For annexin V binding, cells were resuspended in 200 μl of annexin V binding buffer (10 mM HEPES, 150 mM NaCl, 5 mM KCl, 1 mM MgCl_2_, and 1.8 mM CaCl_2_) supplemented with 2 μl of annexin V-FITC (Caltag Laboratories, Burlingame, CA). Annexin-V-positive cells were quantitated by flow cytometry.

### Immunoblotting

Cell lysates were resolved by SDS-PAGE. The proteins were transferred to nitrocellulose (Thermo Fisher) and immunodetected using appropriate primary and peroxidase-coupled secondary antibodies (Genesee Scientific). Proteins were visualized by West Pico and West Dura chemiluminescence Substrate (Thermo Fisher).

### DNA fiber analysis

Exponentially growing cells were pulse-labeled with 50 mM 5-chlorodeoxyuridine (CldU) for 30 min, washed three times with PBS, treated with 2 mM hydroxyurea (HU) for 2 h, washed three times with PBS, incubated again in fresh medium containing 50 mM 5-iododeoxyuridine (IdU) for 60 min, and then washed three times in PBS. DNA fiber spreads were made by a modification of a procedure described previously.^27^ Briefly, cells labeled with IdU and CldU were mixed with unlabeled cells at a ratio of 1:10, and 2 μl cell suspensions were dropped onto a glass slide and then mixed with 20 μl hypotonic lysis solution (10 mM Tris-HCl [pH 7.4], 2.5 mM MgCl2, 1 mM phenylmethylsulfonyl fluoride [PMSF], and 0.5% Nonidet P-40) for 8 min. Air-dried slides were fixed, washed with 1 × PBS, blocked with 5% bovine serum albumin (BSA) for 15 min, and incubated with primary antibodies against IdU and CldU (rat anti-IdU monoclonal antibody [MAb] [1:150 dilution; Abcam] and mouse anti-CldU MAb [1:150 dilution; BD]) and secondary antibodies (anti-rat Alexa Fluor 488-conjugated [1:150 dilution] and anti-mouse Alexa Fluor 568-conjugated [1:200 dilution] antibodies) for 1 h each. Slides were washed with 1x PBS with 0.1% Triton X-100 and mounted with Vectashield mounting medium without 4’, 6-diamidino-2-phenylindole (DAPI). ImageJ software was used to analyze the DNA fibers. For each data set, about 300 fibers were counted for stalled forks, new origins, or elongated forks.

### Viability assay

*Casp2^+/+^* and *Casp2^−/−^* MEF were plated in a 24-well plate at the indicated densities. After four days colonies were visualized by staining overnight with methylene blue 0.1% w/v in methanol/water 50% v/v at room temperature. Cells were washed in PBS.

### Assay for Chromosomal Aberrations at Metaphase

All three stage-specific chromosomal aberrations were analyzed at metaphase after exposure to IR. G1-type chromosomal aberrations were assessed in cells exposed to 3 Gy of IR and incubated for 9-10 h and metaphases were collected.^28, 29^ S-phase–specific chromosome aberrations were assessed after exponentially growing cells irradiated with 2 Gy of IR. Metaphases were harvested following 4 h of irradiation, and S-phase types of chromosomal aberrations were scored. For G2-specific aberrations, cells were irradiated with 1 Gy and metaphases were collected 1-1.5 h post treatment. Chromosome spreads were prepared after hypotonic treatment of cells, fixed in acetic acid·methanol (1:3), and stained with Giemsa. The categories of G1-type asymmetrical chromosome aberrations that were scored include dicentrics, centric rings, interstitial deletions/acentric rings, and terminal deletions. S-phase chromosome aberrations were assessed by counting both the chromosome and chromatid aberrations, including triradial and quadriradial exchanges per metaphase as previously described.^28, 29^ G2-phase chromosomal aberrations were assessed by counting chromatid breaks and gaps per metaphase as previously described.^28, 29^ Fifty metaphases were scored for each post-irradiation time point.

### Statistical Analysis

Statistical comparisons were performed using two-tailed Student’s t test calculated using Prism 6.0 (Graph Pad) software.

## Results

### Loss of caspase-2 increases cell growth

Loss of caspase-2 has been previously shown to increase cellular proliferation rates.^4, 8^ To confirm, we plated E1A/Ras-transformed litter-matched *Casp2*^+/+^ and *Casp2*^−/−^ mouse embryonic fibroblasts (MEF) at low densities and stained for viable cells after 4 days. *Casp2*^−/−^ MEF consistently exhibited an increase in the number of colonies compared to *Casp2^+/+^* cells (Figure 1A). We compared the growth rate of *Casp2^+/+^* and *Casp2^−/−^* cells and caspase-2-deficient cells demonstrated increased proliferation 72 h following plating (Figure 1B, C). To confirm these results, we used BrdU/7AAD staining to determine the cell cycle profiles of cycling *Casp2^+/+^* and *Casp2^−/−^* MEF and found that *Casp2^−/−^* cells had significantly more cells in S-phase and less cells in G1-phase compared to the *Casp2^+/+^* cells (Figure 1D, E). The increased proportion of S-phase cells suggests more caspase-2-deficient cells are synthesizing DNA, while the reduced G1 cells indicate that a decreased proportion of the *Casp2^−/−^* cells are in a quiescent state compared to *Casp2^+/+^* cells. This is consistent with the higher rate of proliferation we observed for caspase-2 knockout cells. Together, these results indicate that caspase-2 plays a role in limiting uncontrolled cell growth.

**Figure 1.**
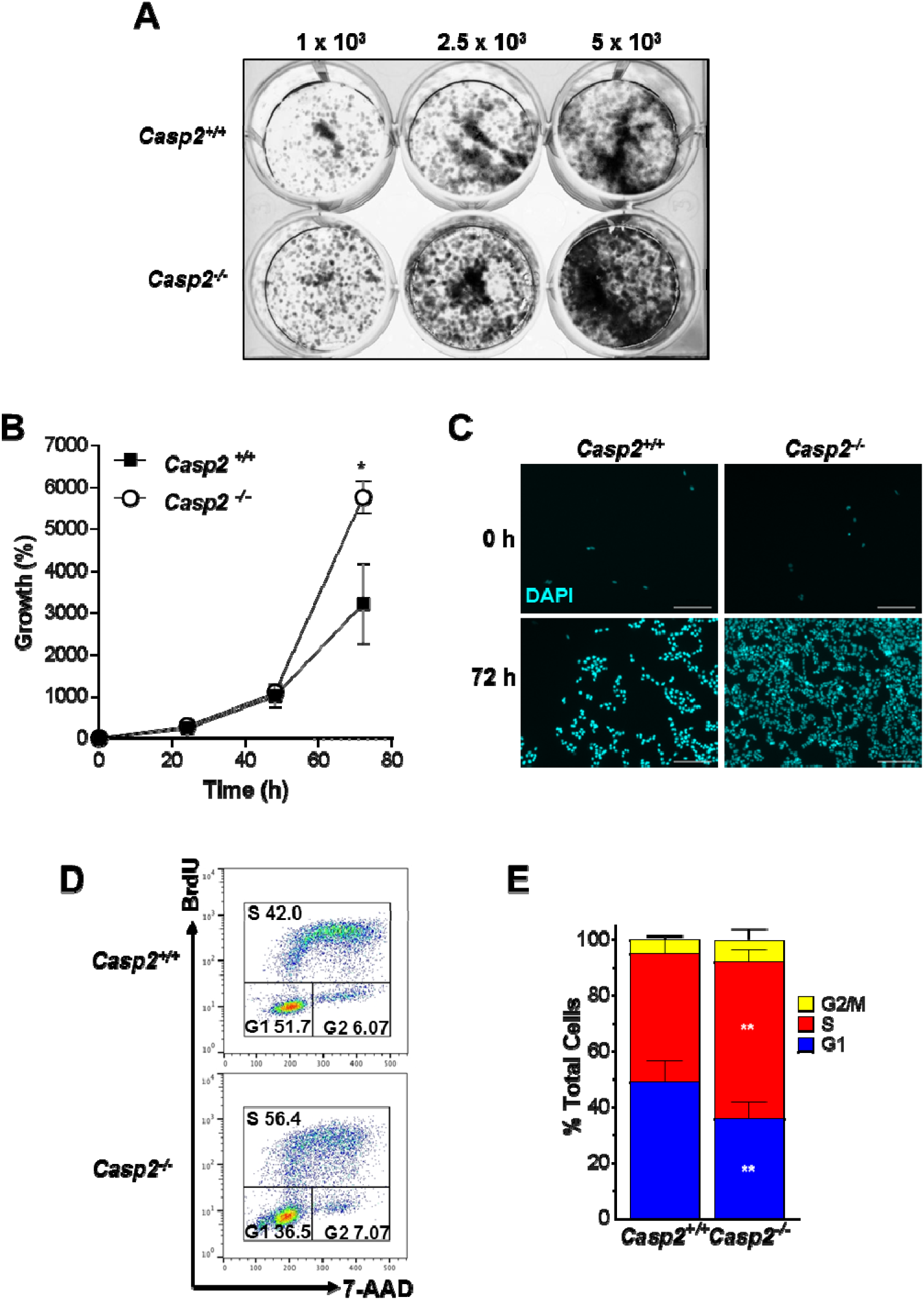
Caspase-2 limits cellular proliferation. **(A)** Litter-matched *Casp2^+/+^* and *Casp2^−/−^* E1A/Ras transformed mouse embryonic fibroblasts (MEF) were plated at the indicated cell number. Viable cells were stained with methylene blue 4 days after plating. Representative images of methylene blue-stained colonies are shown of three independent experiments. **(B)** Litter-matched *Casp2^+/+^* and *Casp2^−/−^* E1A/Ras transformed MEF, plated at low density, were stained with DAPI and imaged at the indicated time points. Total cell number was determined by counting DAPI-positive cells from at least 30 images per well. Results are shown as the percent increase in cell growth compared to 0 h and are the average of three independent experiments plus or minus standard deviation. *p<0.05. **(C)** Representative images are shown from the 0 h and 72 h time points. Nuclei are shown in blue. Bar, 200 μm. **(D)** Untreated, cycling litter-matched *Casp2^+/+^* and *Casp2^−/−^* E1A/Ras MEF were stained with BrdU/7-AAD and analyzed by flow cytometry. Representative flow plots are shown. **(E)** The percentage of cells in G1, S, or G2/M phase for each cell line is shown. Results are the average of seven independent experiments plus or minus standard deviation. **p<0.01.

Because caspase-2 appears to be involved in regulating cell proliferation, we hypothesized that caspase-2 is activated in cells undergoing mitosis. To investigate this, we used the caspase-2 bimolecular fluorescence complementation (BiFC) technique.^19^ This technique relies on the fact that caspase-2 is activated by proximity-induced dimerization following recruitment to its activation platform.^9, 30^ Caspase-2 BiFC uses non-fluorescent fragments of the fluorescent protein Venus fused to the prodomain of caspase-2.^19^ When caspase-2 is recruited to its activation platform the subsequent induced proximity of caspase monomers enforces refolding of the Venus fragments, reconstituting their fluorescence. We used the HeLa.C2 Pro-BiFC line, which stably expresses the caspase-2 BiFC reporter comprising of the prodomain of caspase-2 (aa 1-147) fused to the Venus fragments, Venus N 1-173 (VN) or Venus C 155-249 (VC) separated by the virally derived 2A self-cleaving peptide to ensure equal expression of the BiFC components. ^17^ The C2-Pro BiFC sequence is also linked to an mCherry sequence to permit to visualization of the cells. We used time-lapse confocal microscopy to track caspase-2 BiFC relative to cell division in unstimulated cells. We used the mCherry signal, which fluoresces throughout the cell, to visualize cell division (Figure 2A). We observed caspase-2 BiFC (Venus) around the time of mitosis the majority of cells undergoing division (Figure 2A, 2B, Supplemental Movie 1). In contrast, in cells that did not divide or in cells that died during the course of the time-lapse, the proportion of cells inducing caspase-2 BiFC was equal to the proportion of cells that remained BiFC-negative. This suggests that caspase-2 is activated in dividing cells around the time of mitosis.

**Figure 2.**
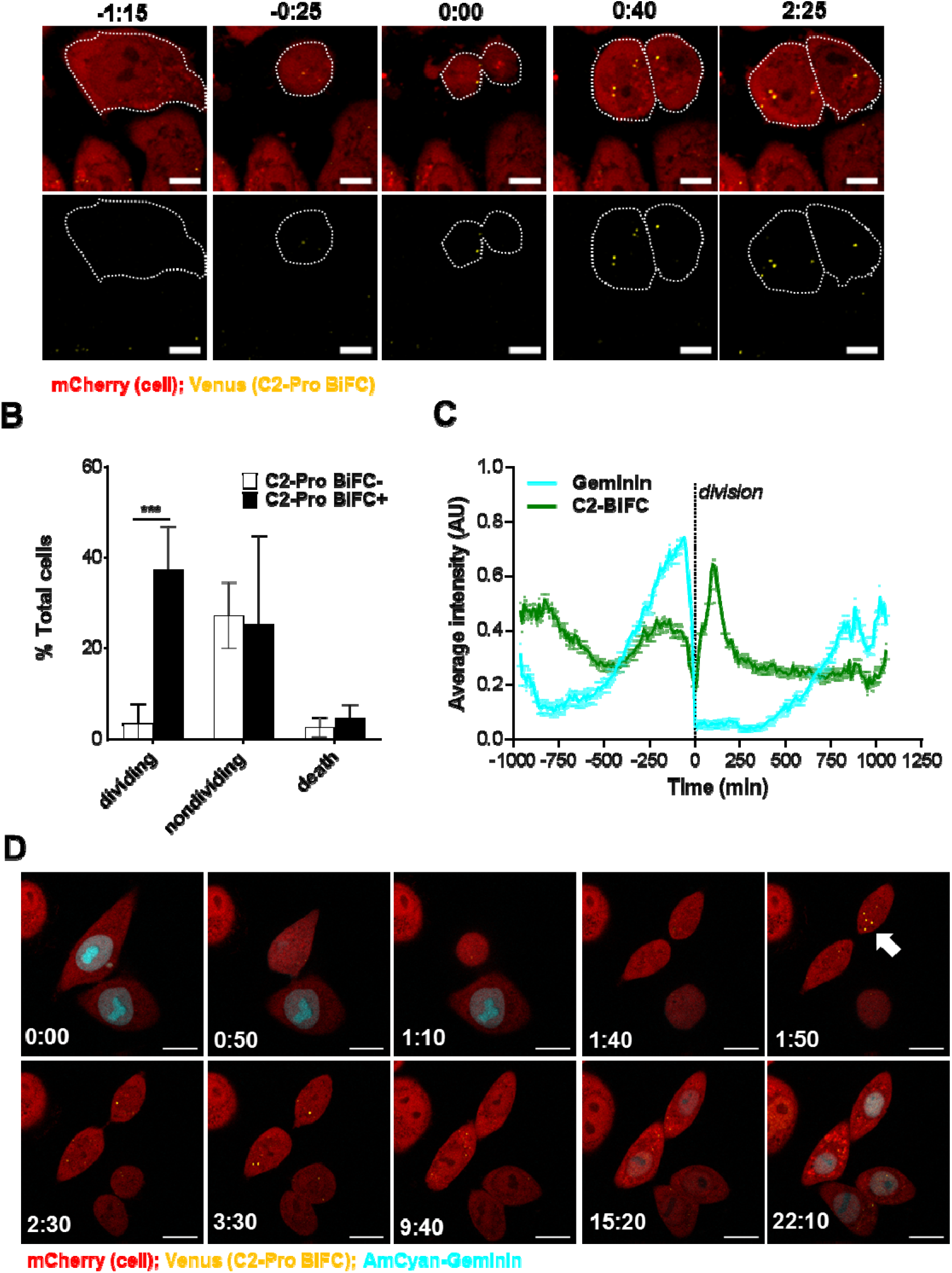
Caspase-2 BiFC is induced in dividing cells. **(A)** HeLa cells stably expressing C2 Pro-VC-2A-C2 Pro-VN-2A-mCherry (HeLa.C2 Pro-BiFC) were imaged every 5 min for 16 h. Representative sequential images show the appearance of caspase-2 BiFC (*green*) in dividing cells (*red*). **(B)** The percentage of mCherry-positive cells that became Venus-positive or remained Venus-negative and divided, did not divide, or underwent cell death was determined from at least 100 cells per experiment. Results are the average of three independent experiments plus or minus standard deviation. ***p<0.001. **(C)** HeLa.C2 Pro-BiFC cells stably expressing AmCyan-Geminin were imaged every 10 min for 24 h. Graph of the dividing cells that became Venus-positive and Cyan-positive is shown. Each point on the Cyan graph (*blue)* is scaled and aligned to each point on the caspase-2 BiFC graph (*green*) that represents the average intensity of Cyan or Venus in the cell at 10 min intervals where time=0 is the time of cell division. The peak of Cyan intensity represens G2-phase of the cell cycle. Error bars represent SEM of 94 individual cell divisions. **(D)** Frames from the time-lapse show representative cells undergoing BiFC (green) relative to geminin expression as measured by the Cyan fluorescence (blue). Scale bars represent 10 μm

To track cell division and the timing of caspase-2 BiFC relative to the cell cycle more accurately, we monitored caspase-2 BiFC relative to a stably expressed fluorescent cell cycle marker, AmCyan-hGeminin. hGeminin is only expressed in G2/M and S-phase of the cycle,^31^ therefore fluorescence of AmCyan-hGeminin is most concentrated in G2 and is degraded in early G1.^32^ When we analyzed the cells that induced caspase-2 BiFC during division, we observed that the peak of caspase-2 fluorescence came 160 min after the peak of the AmCyan-hGeminin signal (Figure 2C, 2D, Supplemental Movie 2). This indicates that caspase-2 activation does not coincide G2/M phase of the cell cycle, but is activated in G1 and early S-phase.

### Loss of caspase-2 results in S-phase specific chromosomal aberrations and delayed exit from S-Phase

To determine which stage of the cell cycle is most impacted by caspase-2 activity, we examined cell cycle-phase specific chromosomal aberrations at metaphase induced by ionizing irradiation (IR) in the presence and absence of caspase-2. G1-specific chromosome aberrations were analyzed in cells treated with 3 Gy and metaphases that were collected 10-12 h post irradiation. S-specific chromosome aberrations were analyzed in cells treated with 2 Gy and metaphases were collected 4-6 h post irradiation and G2-specific chromosome aberrations were analyzed in cells treated with 1 Gy and metaphases were collected 1-1.5 h post irradiation. The frequency of G1-type chromosomal aberrations (mostly of the chromosomal type with frequent dicentric chromosomes)^33^ and G2-type chromosomal aberrations (chromosomal and chromatids) was similar for *Casp2*^+/+^ and *Casp2*^−/−^ cells (Figure 3A). In contrast, S-type aberrations were specifically increased in *Casp2^−/−^* cells. These aberrations consisted primarily of breaks and radials (Figure 3B).

**Figure 3.**
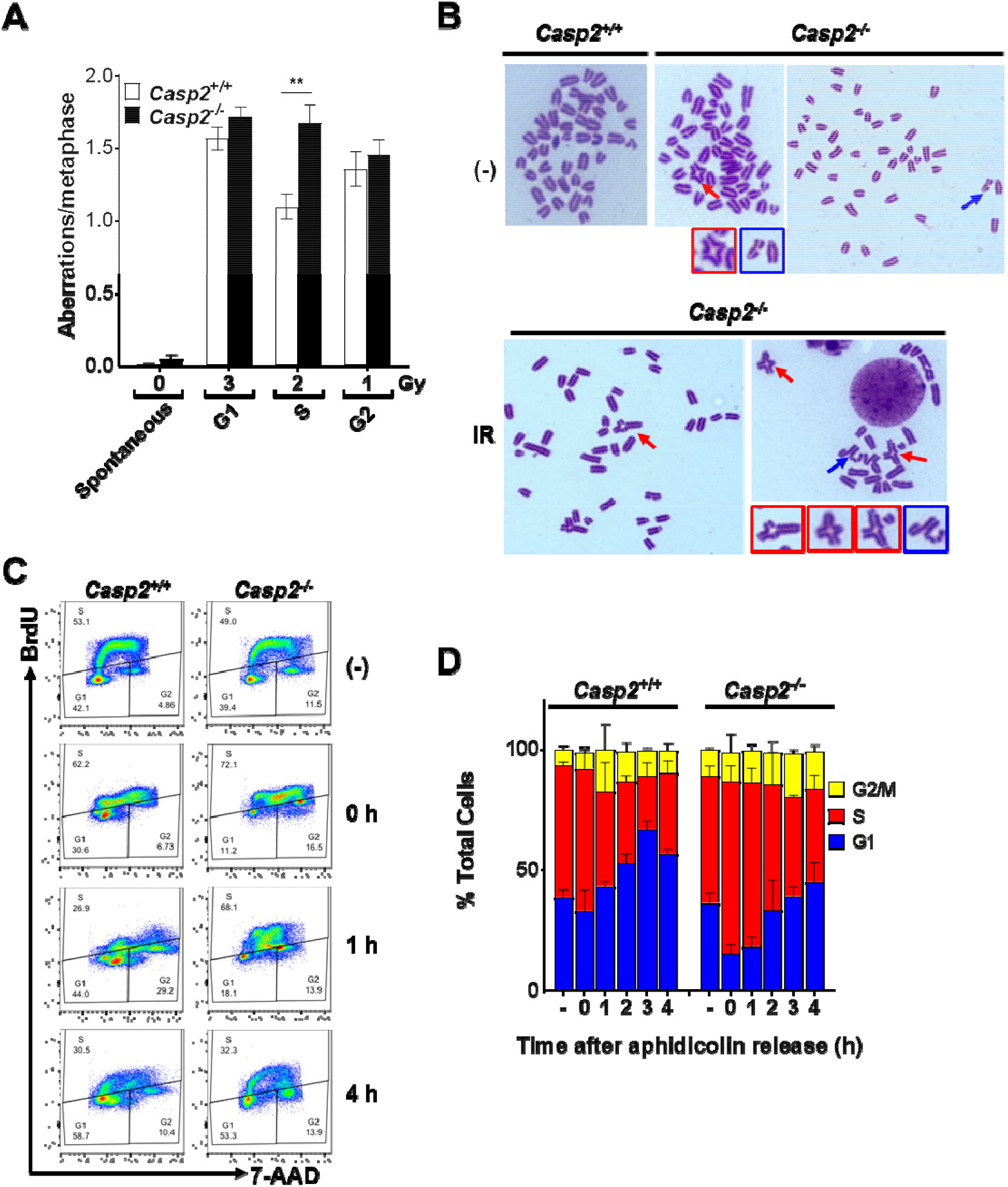
Loss of caspase-2 results in in S-phase specific chromosomal aberrations and delayed exit from S-Phase. **(A)** Litter-matched *Casp2^+/+^* and *Casp2^−/−^* E1A/Ras transformed MEF were treated with the indicated doses of γ-irradiation. Metaphase spreads were prepared and analyzed for chromosomal aberrations that included chromosome and chromatid type breaks and fusions. A total of 35 metaphases were analyzed from each sample and the experiment was repeated three times. **p<0.01. **(B)** Representative images of metaphase spread of untreated (-) or irradiated *Casp2^+/+^* and *Casp2^−/−^* MEF. Red arrows show breaks; blue arrows show exchanges. **(C)** Litter-matched *Casp2^+/+^* and *Casp2^−/−^* E1A/Ras transformed MEF were either left untreated (-) or treated with aphidicolin (1 μM). After 16 h, the treatment was removed and replaced with fresh media. Cells were harvested following a 30 min BrdU (10 μM) pulse at the indicated time points after aphidicolin release and stained with BrdU-FITC/7-AAD. The proportion of cells in S-, G1- and G2-phase was determined by flow cytometry. Representative flow plots are shown. **(D)** The percent of cells in each phase of the cell cycle was determined for each time point following aphidicolin release. Results are the average of 3 independent experiments plus or minus standard deviation.

The specific increase in S-phase specific chromosomal aberrations in the absence of caspase-2 suggests a role for caspase-2 in ensuring normal S-phase progression. Therefore, we investigated the impact of loss of caspase-2 on cell cycle recovery after S-phase arrest. We treated *Casp2*^+/+^ and *Casp2*^−/−^ MEF with aphidicolin for 16 h to synchronize the cells in S-phase. Immediately after release from S-phase arrest (0 h), there was a greater proportion of *Casp2*^−/−^ MEF in S-phase compared to *Casp2*^+/+^ MEF. In addition, over four hours following release from arrest the proportion of *Casp2*^+/+^ MEF in S-phase reduced as the cells moved into G2 and then G1. In contrast, *Casp2*^−/−^ cells exhibited a delay in exit from S-phase and concomitant entry into G1 (Figure 3C, D). This suggests that the exit from S-phase was delayed in *Casp2*^−/−^ MEF.

### Caspase-2 protects from stalled replication forks and DNA damage

The delayed recovery from S-phase arrest we observed in *Casp2*^−/−^ cells could be due to DNA replication stress. Therefore, we investigated the role of caspase-2 in replication fork dynamics following transient genotoxic stress-induced replication blockage. We used a DNA fiber assay to evaluate restart and recovery of replication forks after hydroxyurea (HU) treatment. HU induces replication fork stalling and S-phase arrest by depleting the available nucleotide pool for DNA polymerases.^34^ Cells were pulse-labeled with 5-iododeoxyuridine (IdU), treated with HU for 2 h to induce replicative stress, and then washed and pulse-labeled with 5-chlorodeoxyuridine (CldU). Individual DNA fibers, which incorporated the CldU and/or IdU pulses, were detected with fluorescent antibodies against those analogs. We noted three types of DNA fiber tracts: those with ongoing elongation forks (CIdU+IdU), stalled forks (CIdU), and newly initiated forks (IdU) representing new origins of DNA replication (Figure 4A-F). Caspase-2-deficient MEF demonstrated a significantly higher frequency of stalled replication forks (Figure 4B), new origins of replication (Figure 4C), and a reduced frequency of CldU+IdU forks, indicating ongoing replication (Figure 4D). This suggests that the loss of caspase-2 prevents or delays reinitiation of stalled replication forks that can result in S-phase arrest, while also resulting in excessive replication-origin firing. When replication is stalled, the length of DNA tract is impacted.^35^ Therefore, to assess the impact of caspase-2 loss on DNA tract length, we measure the length of the total tract of CldU + IdU and the length of CldU or IdU labeled DNA fibers in *Casp2^+/+^* and *Casp2*^−/−^ cells. As expected, the length of the CldU fibers, which represents the DNA prior to replication stress, were the same across the cell types. The length of IdU was significantly reduced in *Casp2*^−/−^ cells compared to wild type (Figure 4E, F). This reduced fiber length indicates a reduced DNA fork speed and, thus, increased replication stress in the absence of caspase-2. Thus caspase-2 promotes replication fork progression.

**Figure 4.**
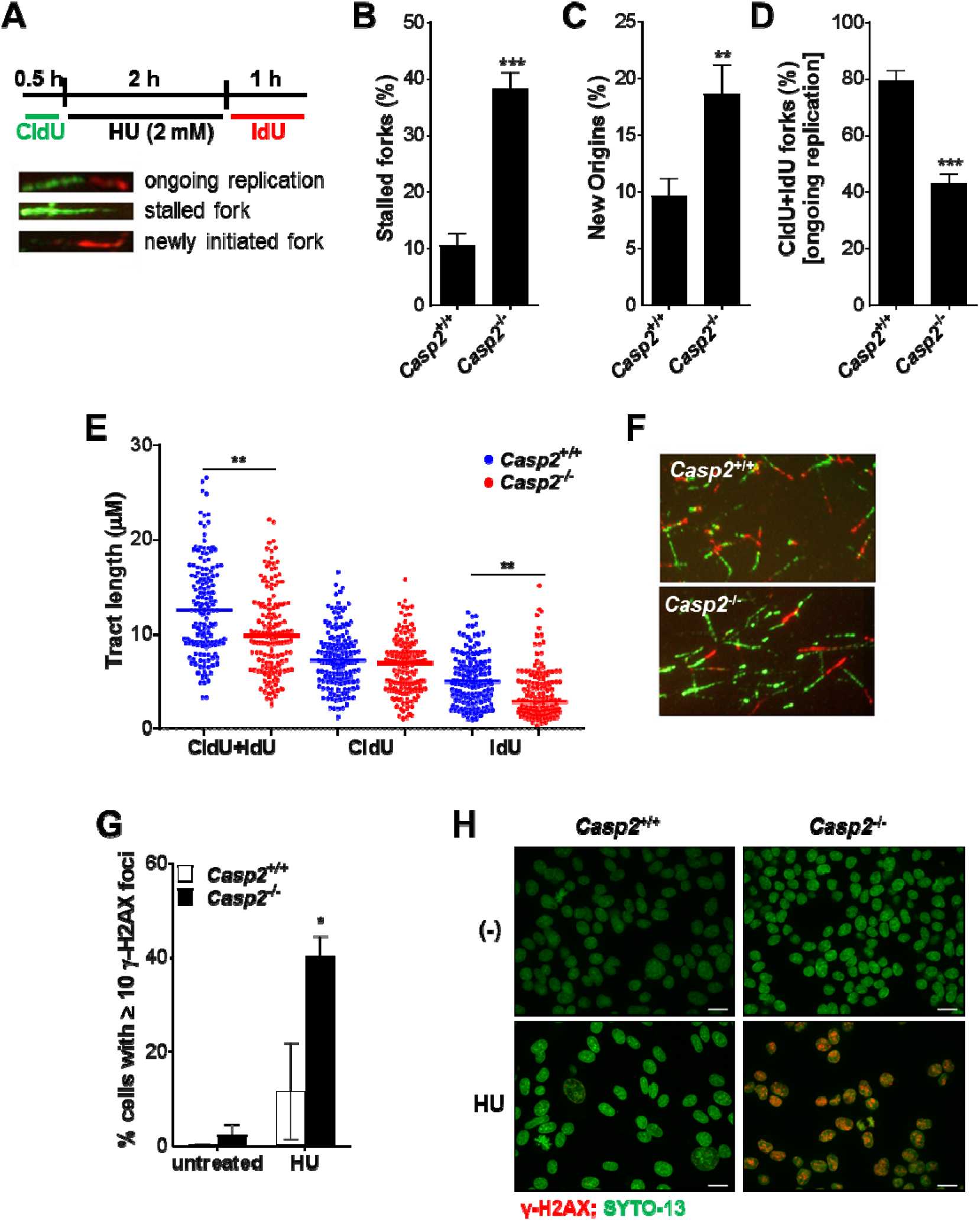
Loss of caspase-2 is associated with stalled replication forks and increased DNA damage. **(A)** Scheme for dual labeling of DNA fibers to evaluate replication restart or recovery following HU-induced replication fork stalling. **(B to E).** Quantitative analysis of the DNA fiber replication restart assay after HU treatment shows the percentage of stalled forks **(B)**, the percentage of new origins **(C)**, total tracts with both CldU and IdU labels at the fork (CldU+IdU) **(D)**, and tract lengths **(E)** in *Casp2^+/+^* and *Casp2^−/−^* MEF. The experiment was performed three times, and for each condition about 300 fibers were analyzed. Error bars represent the standard deviation from three independent experiments. **p<0.01, ***p<0.001. **(F)** Representative images of replication tracts from litter-matched *Casp2^+/+^* and *Casp2^−/−^* E1A/Ras - transformed MEF after HU treatment showing CIdU (5-chloro-2’-deoxyuridine, green) and IdU (5-iodo-2-deoxyuridine, red) labeled tracts are shown. **(G)***Casp2*^+/+^ and *Casp2*^−/−^ MEF treated with DMSO or HU (2 mM) for 2 h followed by recovery for 24 h were stained for γ-H2AX. γ-H2AX foci were counted per cell and the percentage of cells with ≥ 10 foci/cell was calculated from at least 30 images per treatment. Results are the average of three independent experiments plus or minus standard deviation. *p<0.05. **(H)** Representative images from (G) are shown as maximum intensity projections of 5 image Z stacks, with nuclei shown in green and γH2AX foci in red. Bar, 50 μm.

Excessive replication fork stalling induced by HU treatment can lead to fork collapse and breakage in the form of DNA double strand breaks (DSBs).^36^ To probe the effects of loss of caspase-2 on DSBs induced by replication stress, we stained for phosphorylated H2AX (γ-H2AX). γ-H2AX forms foci at DSBs and therefore is a reliable and sensitive indicator of DSBs.^37^ We treated *Casp2*^+/+^ and *Casp2*^−/−^ MEF with HU for 2 h and quantitated the percentage of cells with γH2AX foci 24 h later. We observed a significantly higher level of cells with γH2AX foci in *Casp2*^−/−^ MEF compared to the wild type MEF (Figure 4G, H). These data indicate a higher level of DNA damage following replication stress in the absence of caspase-2. This suggests that caspase-2 may function to prevent DNA damage or to facilitate DNA repair.

### Loss of caspase-2 is not associated with impaired activation of known cell cycle checkpoints

Stalled replication forks lead to activation of ATR and its substrate Chk1.^38^ Chk1 triggers a G2/M checkpoint by inhibiting Cdc25C-mediated activation of Cyclin B^39^ and also triggers an intra S-phase checkpoint through inhibition of Cdc25A-mediated activation of Cyclin A/E.^40^ Given the increase in stalled forks and delayed exit from S-phase in the absence of caspase-2, we measured the impact of loss of caspase-2 on ATR and Chk1 activation. We measured checkpoint activation in *Casp2*^+/+^ and *Casp2*^−/−^ MEF treated with HU (Figure 5A). Chk1 was phosphorylated immediately after HU release to a similar extent with and without caspase-2 and minimal differences in Chk1 phosphorylation between the two cell lines were noted over time. Interestingly, phosphorylation of ATR increased 2 h following HU release in *Casp2*^+/+^ MEF and this was inhibited in *Casp2*^−/−^ MEF. However, the phosphorylation that was detected was at Serine 428. Phosphorylation of ATR at this site is not required for Chk1 phosphorylation and therefore not associated with ATR activation.^41^ Because antibodies for the phospho-ATR site that is associated with activation (threonine 1929) are not available for murine ATR, we used CRISPR/Cas9 to generate human caspase-2-deficient U2OS cells to determine if caspase-2 is required for ATR activation. As in the MEF, loss of caspase-2 had no effect on Chk1 phosphorylation induced by two different doses of HU or the DNA damage inducers: etoposide, camptothecin or topotecan (Figure 5B). Phosphorylation of ATR at T1989 was not substantially impacted by the loss of caspase-2 under any of the treatment conditions. We similarly generated caspase-2-deficient HeLa cells using CRISPR/Cas9. Similar to the U2OS cells, no difference in Chk1 phosphorylation was observed (Figure 5C). Finally, we examined Chk1 phosphorylation in response to S-phase arrest induced by aphidicolin in U2OS cells (Figure 5D). Following release from aphidicolin, we did not detect any difference in Chk1 phosphorylation.

**Figure 5.**
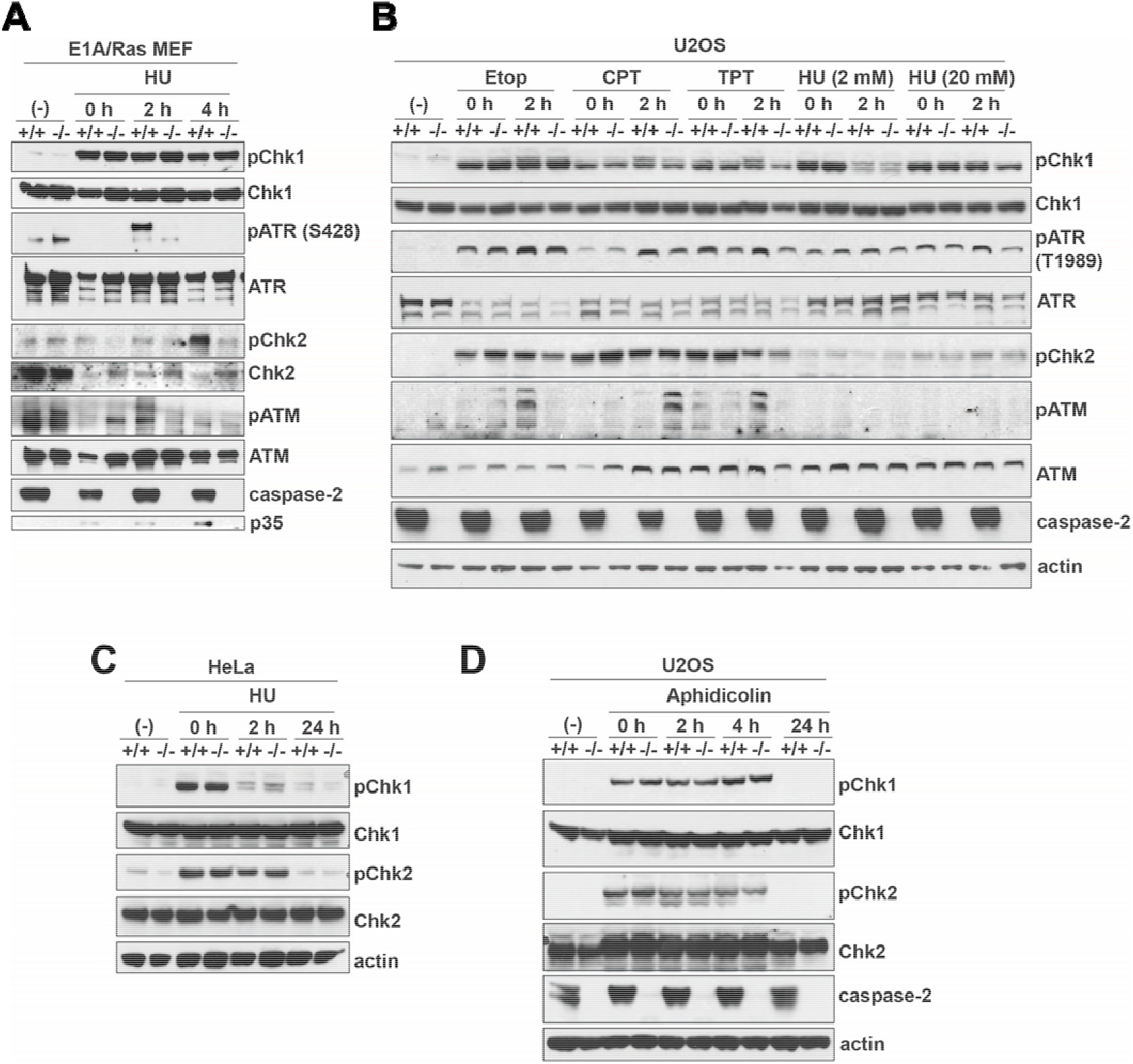
Loss of caspase-2 does not impair ATR or ATM checkpoints. **(A)** Litter-matched *Casp2^+/+^* and *Casp2^−/−^* E1A/Ras-transformed MEF were either left untreated (-) or treated with hydroxyurea (HU, 2 mM) for 2 h followed by replacement with fresh media. Cells were harvested at the indicated time points following wash-out of HU. Cell lysates were immunoblotted for the indicated checkpoint proteins and their phosphorylated counterparts. Actin was used as a loading control. **(B)** U2OS cells or CRISPR/Cas9-generated caspase-2-deficient U2OS cells were left untreated or treated with etoposide (20 μM), camptothecin (100 μM), topotecan (100 μM) for 4 h or with HU (2 mM or 20 mM) for 2 h followed by replacement with fresh media. Cells were harvested at the indicated time points following wash-out of the drug and lysates were immunoblotted for the indicated proteins. **(C)** HeLa cells or CRISPR/Cas9-generated caspase-2-deficient Hela cells were treated as in (A). Lysates were immunoblotted for the indicated proteins. **(D)** Caspase-2 wild type or deficient U2OS cells were treated with aphidicolin (1 μM) for 16 h followed by replacement with fresh media. Cells were harvested at the indicated times and lysates were immunoblotted for the indicated proteins. Each experiment is representative of 2-6 independent experiments.

ATM regulates the intra-S phase checkpoint through Chk2 activation.^42^ Similar to ATR, we saw some induction of ATM phosphorylation in MEF 2 h following HU release that was diminished in the caspase-2 knockout cells (Figure 5A). However, this was not accompanied by phosphorylation of the ATM substrate, Chk2.^43^ In MEF, Chk2 phosphorylation was detected at the later 4 h timepoint and this appeared caspase-2-dependent. ATM is activated primarily by double strand breaks,^44^ therefore, the later Chk2 activation may be a result of DSBs resulting from collapsed stalled replication forks. To determine the effect of loss of caspase-2 on ATM activation in response to DSBs, we measured ATM and Chk2 phosphorylation in response to etoposide compared to the single strand DNA break inducers, camptothecin and topotecan, and to HU (Figure 5B). We observed strong phosphorylation of Chk2 immediately after release from etoposide, camptothecin, or topotecan in the presence and absence of caspase-2. In contrast, HU did not induce much Chk2 phosphorylation. ATM phosphorylation was detected at 2 h post release in response to etoposide, camptothecin, or topotecan and this was diminished in caspase-2 knockout cells. ATM phosphorylation was not induced by HU in the U2OS cells. In HeLa cells and in U2OS cells, we detected strong Chk2 phosphorylation in response to HU and aphidicolin respectively in the presence and absence of caspase-2 (Figure 5C and D). Taken together, these results suggest that caspase-2 is either activated downstream of ATM and ATR or in parallel to these checkpoints.

### Caspase-2 induced cell cycle regulation is independent from its ability to induce apoptosis

Caspase-2 induces apoptosis through cleavage of the pro-apoptotic protein BID and subsequent permeabilization of the outer mitochondrial membrane, cytochrome c release, and downstream caspase activation.^45, 46^ However, it is unclear if the pathway engaged by caspase-2 to regulate the cell cycle is the same pathway that induces apoptosis. To investigate this, we overexpressed Bcl-X_L_ in the *Casp2*^+/+^ and *Casp2*^−/−^ MEF. Bcl-X_L_ potently blocks caspase-2 induced apoptosis,^19^ and these cells were completely resistant to the apoptosis inducers actinomycin D, staurosporine and etoposide (Figure 6A). Nevertheless, the cell cycle profile of the Bcl-X_L_-overexpressing *Casp2*^+/+^ cells was unchanged compared to that of the vector-transduced *Casp2*^+/+^ MEF (Figure 6B). This suggests that the association between the increased frequency of S-phase defects and loss of caspase-2 is not due to apoptosis induced by caspase-2, indicating that a distinct pathway is engaged to induce the observed cell cycle-related effects. To explore this further, we treated *Casp2*^+/+^ and *Casp2*^−/−^ MEF expressing vector or Bcl-X_L_ with camptothecin for four hours and measured both cell cycle arrest at 4 h (Figure 6C) and apoptosis at 24 h (Figure 6D). When we challenged the cells with low doses of camptothecin, *Casp2*^+/+^ cells accumulated in S-phase and this was increased in *Casp2*^−/−^ cells. At higher doses of camptothecin, *Casp2*^+/+^ cells accumulated more in G2/M-phase. This was slightly increased in the absence of caspase-2 but the effect was variable. The proportions of G1 cells steadily decreased with increasing dose of camptothecin. The increases in S-phase arrest at lower doses, G2-arrest at high doses and decreased G1 cells was not changed by the overexpression of Bcl-X_L_ (Figure 6C). These treatment conditions induced minimal apoptosis after 24 h that was potently blocked by overexpression of Bcl-X_L_ (Figure 6D). Together these results indicate that the S-phase defects associated with caspase-2-deficiency are independent of the ability of caspase-2 to induce apoptosis.

**Figure 6.**
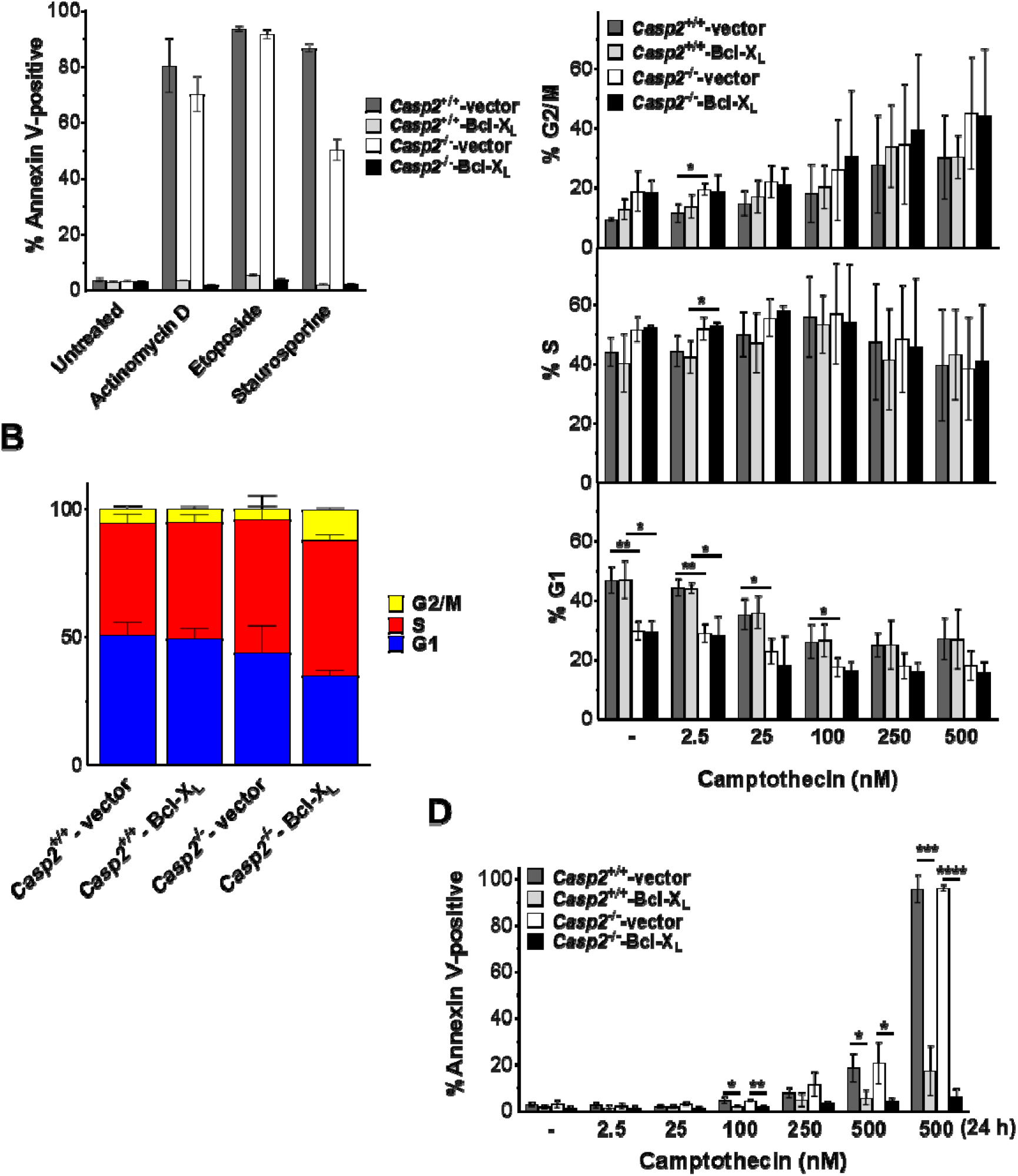
The role of caspase-2 in cell division is independent of its ability to induce apoptosis. **(A)** Litter-matched *Casp2^+/+^* and *Casp2^−/−^* E1A/Ras transformed MEF stably expressing vector or Bcl-X_L_ were treated with actinomycin D (0.5 μM), etoposide (50 μM), or staurosporine (1μM) for 24 h. Apoptosis was measured by Annexin V staining. Cycling *Casp2^+/+^* and *Casp2^−/−^* MEF stably expressing vector or Bcl-X_L_ were untreated **(B)** or were treated with the indicated doses of camptothecin for 4 h **(C)**. Cells were harvested following a 30 min BrdU (10 μM) pulse and stained with BrdU/7-AAD to determine the percentage of cells in G1, S or G2/M-phase of the cell cycle. Results are an average of 3 independent experiments plus or minus standard deviation. (**D**) *Casp2^+/+^* and *Casp2^−/−^* MEF stably expressing vector or Bcl-X_L_ were treated with the indicated doses of camptothecin for 4 h or 24 h. Apoptosis was measured at 24 h by Annexin V staining. Results are the average of 3 independent experiments plus or minus standard deviation.

## Discussion

Here we report that caspase-2 is activated during mitosis and loss of caspase-2 is associated with S-phase specific defects, including DNA fork stalling, delayed exit from S-phase, and S-phase specific chromosomal aberrations. These functions appear to be independent of caspase-2’s ability to induce apoptosis and loss of caspase-2 has minimal impact on the intra S-phase checkpoint. Altogether, our results demonstrate that caspase-2 plays a key non-apoptotic role in the regulation of DNA replication that protects against genomic instability.

Our data indicate that caspase-2 is activated during mitosis. Notably, this activation occured during G1 and early S-phase of the cycle. The activation of caspase-2 in G1 or early S is consistent with the S-phase associated defects. The higher rate of S-phase arrest in *Casp2^−/−^* cells may suggest that the activation of caspase-2 in G1 leads to protection of the G1/S transition. Caspase-2 is also required for timely S-phase progression. Following S-phase arrest, cells progress through S-phase to G2 and G1 more quickly than when caspase-2 is absent. This indicates that in the absence of caspase-2, cells struggle to exit S-phase. Thus, our data indicates that caspase-2 has an important role in G1/S phase of the cell cycle. During S-phase of the cell cycle, the genome of the cell is duplicated.^47^ Any errors sustained during this process can manifest as replication stress and can result in stalled or collapsed replication forks.^48^ Our data show that caspase-2 protects against stalled replication forks and against excessive replication-origin firing induced by replication stress. Stalled replication forks produce single strand DNA that, when collapsed, can be converted to DSBs, the accumulation of which promotes genomic instability.^49^ Excessive replication origin firing can lead to the depletion of necessary nutrients and metabolites required for correct genome replication, again contributing to genomic instability.^50, 51^ Several groups, including our own, have provided evidence that caspase-2 protects against genomic instability.^5, 7, 52, 53^ Loss of caspase-2 has been shown to be associated with higher levels of aneuploidy,^53, 54^ polyploidy,^52^ and genome duplication.^7^ Our data provides key evidence that the mechanism by which caspase-2 that protects against genomic instability is by protecting DNA replication forks during S-phase of the cell cycle.

The primary S-phase-associated cell cycle checkpoint is the intra-S-phase checkpoint that is required to ensure genomic integrity and to prevent errors during DNA replication.^55^ ATR is a critical mediator of the intra-S-phase checkpoint and is activated by ssDNA, DSBs, and it also regulates origin firing in normal S-phase.^56^ Several studies indicate that the complex between replication protein A and ssDNA is the convergence point for different types of lesions to activate ATR.^57, 58^ In contrast, DSBs activate ATM.^44, 59^ Surprisingly, although loss of caspase-2 was associated with increased stalled replication forms and origin firing, it had minimal effects on ATR or Chk1 phosphorylation. This would suggest that caspase-2 functions downstream of, or independently of, ATR and Chk1 in response to stalled replication forks. Chk1 has been proposed as an inhibitor of caspase-2 in response to IR-induced DNA damage.^15^ Inhibition of Chk1 potentiates caspase-2 dependent apoptosis by derepressing ATM-mediated phosphorylation of the upstream caspase-2 activator PIDD. However, this appears to be specific to the IR pathway since inhibition of Chk1 alone is not sufficient to induce caspase-2.^60^ Therefore, it is unclear if this phenomenon could impact the response to replication stress. The observed decrease in ATM autophosphorylation in the absence of caspase-2 suggests impaired ATM activation that could also contribute to the intra S-phase checkpoint.^61^ However, this decrease was not accompanied by a decrease in activation of its substrate Chk2. ATM phosphorylates additional cell cycle proteins including p53,^62^ BRCA1^63^ and NBS1^64^ and it is possible that loss of caspase-2 has an impact on a different ATM substrate to impact this or other checkpoints. In addition, it has been demonstrated that Chk2 can be activated independently of ATM in response to HU treatment.^65–67^ The fact that loss of caspase-2 potentiates tumorigenesis and genomic instability in an ATM-deficient background^68^ argues against any interdependence of these two pathways and suggests rather that caspase-2 and ATM function in parallel to protect against genomic instability.

During cell division, caspase-2 is subject to two separate phosphorylation events that are reported to attenuate its activation downstream of activation platform assembly. These phosphorylation events are induced by Cdk-cyclin B1 and Aurora B kinase at S340 and S384 respectively.^69, 70^ Cdk-cyclin B1 is required for G2/M transition,^71^ while Aurora B kinase is a spindle checkpoint protein and is essential for the segregation of chromosomes.^72^ It has been proposed that these inhibitory phosphorylation events are to prevent caspase-2 from inducing apoptosis during cell division, a process referred to as mitotic catastrophe.^69, 70^ However, we noted that caspase-2 activation in dividing cells was primarily in G1 phase rather than G2, indicating that this is a separate event. In addition, while we noted a strong association between caspase-2 activation and cell division, we did not observe a similar association with cell death. This demonstrates that apoptosis is not the primary outcome of this caspase-2 activation during cell division. Therefore, it is possible that an alternative function of these phosphorylation events is to fine tune caspase-2 activation during different stages of the cell cycle, allowing its activation during G1- and S-phase and downregulating its activity during G2/M.

Treatment with inducers of G2/M arrest including nocodazaole and Plk1 inhibition has been shown to induce caspase-2-dependent apoptosis as a mechanism to remove aneuploidy cells.^54^ However, it is not clear whether caspase-2 regulates cell cycle during the DNA damage response through activation of the same pathway components that are involved in caspase-2-dependent apoptosis or if caspase-2-mediated cell cycle regulation is independent of its ability to induce apoptosis. Bcl-X_L_ is an effective inhibitor of caspase-2 –induced apoptosis via its ability to block the caspase-2 substrate BID and to inhibit mitochondrial outer membrane permeabilization.^73^ Our evidence indicates that the cell cycle effects associated with the loss of caspase-2 is not phenocopied by Bcl-X_L_ overexpression. This indicates that the cell cycle functions of caspase-2 leading to DNA fork protection and prevention of DNA damage are not simply due to removal of damaged cells by caspase-2-dependent apoptosis. Our evidence further suggests that the regulation of cell cycle by capsase-2 is mechanistically independent of its ability to induce apoptosis. Caspase-2 has been shown to cleave MDM2 to increase p53 stabilization in response to Aurora B kinase inhibition.^11^ This has been shown to be important for inducing cell cycle arrest in response to cytokinesis failure.^52^ It is possible that cleavage of MDM2 rather than BID is responsible for protecting DNA replication forks. However, it remains possible that these effects are due to an, as yet, unidentified caspase-2 target.

Another group showed that PIDD-deficiency is associated with decreased recovery from stalled DNA replication forks induced by UV.^74^ The similarity between the results of the latter study and our results may suggest that the PIDDosome is responsible for the function of caspase-2 in protecting replication forks. We identified the nucleolar PIDDosome comprising of nucleophosmin (NPM1), PIDD, and RAIDD as a major caspase-2 activating complex in response to topoisomerase I inhibitors that produce ssDNA.^75^ We showed that inhibition of NPM1 overcomes caspase-2 associated growth arrest^75^. This may suggest that the nucleolar PIDDosome drives the cell cycle events we report here. However, Lin et al concluded that PIDD facilitates DNA-PKcs binding to ATR and, in contrast to our results, showed that the ATR pathway was compromised in the absence of PIDD.^74^ PIDD has also been reported to bind PCNA to facilitate trans-lesion synthesis following UV-induced damage.^76^ This complex was shown to be independent of caspase-2 function. This may suggest that PIDD is acting independently of caspase-2 to activate ATR, but the nucleolar PIDDosome is required for efficient resolution of stalled forks and prevention of DSBs. To fully understand the conditions that lead to caspase-2 activation during cell cycle and how this leads to DNA replication fork protection, further exploration of the complexes that lead activation platform in this process will be essential.

In conclusion, our data indicate an active role for caspase-2 in ensuring correct cell cycling that does not simply lead to removal of damaged cells by apoptosis and that this pathway does not overlap with caspase-2’s pro-apoptotic function. Importantly, our results provide strong evidence that caspase-2 engages multiple cellular functions to safeguard against genomic instability and tumor progression.

## Acknowledgements

We would like to thank Jennifer Martinez (NIEHS) for careful reading of this manuscript. Funding for this project includes NIH/NIGMS R01GM121389 and NIH/NCI R21CA256606 (LBH). This project was supported by the Cytometry and Cell Sorting Core at Baylor College of Medicine with funding from the NIH (P30 AI036211, P30 CA125123, and S10 RR024574) and the expert assistance of J. M. Sederstrom. We would like to acknowledge the Texas Children’s Hospital William T. Shearer Center for Human Immunobiology for their generous support for this research and the expert assistance of Rebecca Kairis.

